# Effects of Aversive Classical Conditioning on Pupil Dilation and Microsaccades

**DOI:** 10.1101/2022.06.09.495508

**Authors:** W.M. Friedl, Andreas Keil

## Abstract

Unintentional shifts of gaze and the modulation of pupil size are both highly automated processes that serve to regulate the initial influx of visual information. The present work investigated these mechanisms as they came to differentially respond to initially unthreatening stimuli following aversive conditioning. The classical conditioning experimental paradigm employed simple grating stimuli (Gabor patches) shown individually at various (5) on-screen locations, one of which was paired with a noxious auditory unconditioned stimulus (US). Aversively paired Gabor patches elicited an attenuated initial constriction of the pupil (pupillary light response; PLR) along with more rapid re-dilation compared to otherwise identical but unpaired gratings. Modulation of both the PLR and the rate of re-dilation following conditioning showed pronounced individual differences. Rapid eye movements away from fixation (microsaccades) were suppressed for the aversively associated compared to unassociated stimulus locations at several post-stimulus latencies. Mutual information between pupil dilation and microsaccade rate, meanwhile, did not differ between the aversively associated and unassociated visual on-screen locations. Together, these results suggest that measures of pupil diameter and microsaccade rate supply complementary information on early-stage processes of associative learning through experience.

---

The ability to learn visual cues that foretell dangerous environments is an evolutionarily powerful advantage afforded to sighted organisms. Classical (Pavlovian) conditioning studies have detailed a canonical “fear circuit” originating in the amygdala (e.g Blanchard & Blanchard, 1972; LeDoux, Cicchetti, Xagoraris, & Romanski, 1990; Thompson, 1983; reviewed in LeDoux, 2000) often active when learning threatening cues. However, how such learning impacts sensory systems—particularly the visual system of human subjects—is still unclear. At the gateway of the visual system, eye movements and pupil dilation regulate the influx of information: Altering either of these behaviors will, therefore, affect all downstream processing stages. The present study investigated how small eye movements during attempted fixation (microsaccades) and pupil diameter responded when subjects viewed differentially conditioned on-screen locations.

Fixational eye movements (including microsaccades) and pupillary dilation/constriction both facilitate the active extraction of important information from the visual environment (Mathôt, 2018; Najemnik & Geisler, 2005). Fast, abrupt, and (relatively) large eye movements made during free visual exploration (saccades) focus high-acuity retinal processing resources on points throughout the visual field (Liversedge & Findlay, 2000; Whittaker & Cummings, 1990). Smaller velocity saccades made while trying to maintain stationary gaze (fixational or microsaccades) are now thought to largely share the structural and functional properties of larger saccades (Hafed & Krauzlis, 2012; Martinez-Conde, Macknik, & Hubel, 2004; Otero-Millan, Macknik, Serrano-Pedraza, & Martinez-Conde, 2008). Pupil responses reflect a balance between parasympathetic (constriction) and sympathetic (dilation) tone (Bradley, Sapigao, & Lang, 2017; Hudspeth et al., 2013, p. 1030). The primary function of the pupil is to optimize visual acuity under varying luminance conditions (Campbell & Gregory, 1960; Laughlin, 1992), but pupil diameter is also impacted by cognitive factors such as anticipation and arousal (reviewed in Peinkhofer, Knudsen, Moretti, & Kondziella, 2019 and in Sirois & Brisson, 2014). Both microsaccades (Carrasco, 2018; Engbert & Kliegl, 2003; Hafed & Clark, 2002; Laubrock, Engbert, Rolfs, & Kliegl, 2007; Yuval-Greenberg, Merriam, & Heeger, 2014; but see Horowitz, Fine, Fencsik, Yurgenson, & Wolfe, 2007) and pupillary responses (Binda, Pereverzeva, & Murray, 2013; Mathôt, Van der Linden, Grainger, & Vitu, 2013; Naber, Alvarez, & Nakayama, 2013) have been suggested to index the deployment of covert attention, and biased processing of aversively conditioned cues has also been attributed, in part, to them preferentially attracting attention (e.g. Bradley, 2009; Dayan, Kakade, & Montague, 2000; Lang, Bradley, & Cuthbert, 1997; Mackintosh, 1975; Panitz, Keil, & Mueller, 2019). Modulation of the pupil and saccadic eye movements also share neural generators (Benedetto & Binda, 2016; Lorber, Zuber, & Stark, 1965) including intermediate layers of the superior colliculus (SCi; Joshi & Gold, 2020; Wang & Munoz, 2015; 2021).

Given their shared functions and neural architecture, interactions between microsaccade rate and pupil responses are increasingly being investigated (e.g. Mathôt, Van der Linden, Grainger, & Vitu, 2013; Pandey & Ray, 2021; Privitera, Carney, Klein, & Aguilar, 2014; Strauch, Greiter, & Huckauf, 2018; Wang, Huang, Brien, & Munoz, 2020) but have yet to be considered in the context of aversive conditioning. As distinct dependent measures of human classical conditioning, pupil activity has a long record (Reinhard & Lachnit, 2002; Voigt, 1968), while conditioning effects on microsaccade rate are unknown. A recent meta-analysis of pupil response in classical conditioning studies (Finke, Roesmann, Stalder, & Klucken, 2021) has furnished effect size estimates for many study-level variables (e.g. modality of the conditioned stimulus); the present work complements this by supplying individual-level effect size estimates from a sample of 51 healthy participants. Mutual information was calculated to measure the strength of association between pupil dilation and microsaccads during aversive associative learning.

## Method

Pupil diameter and microsaccade rates were extracted from eye tracker data collected as part of a larger EEG study, results of which have been published elsewhere (Friedl & Keil, 2021). Here we only detail the procedural and data analysis steps applicable to pupillary and eye movement responses.

Participants completed a delayed, differential classical conditioning paradigm. Differential conditioning utilizes at least two distinct conditioned stimuli (here, five), one of which (the CS+) comes to predict the presence of the unconditioned stimulus (US), while the other (the CS-) comes to predict the absence of the US (Domjan, 2018). In delayed conditioning, the US occurs after the CS+ has been present for some time in each association trial, and the CS+ and US co-terminate. The auditory US and visual CS+ and CS- stimuli are described below.

### Participants

Fifty-three students at the University of Florida took part in the study for course credit. One participant chose to discontinue testing due to discomfort with the auditory US, and data from one participant were excluded because of a defective EEG sensor net, leaving a final sample of 51 participants (19 males; M_age_ = 19.49 years; SD_age_ = 1.22). All participants reported no personal or family history of epilepsy or photic seizures and normal or corrected-to-normal vision. Participants provided informed consent in accordance with the Declaration of Helsinki and the institutional review board of the University of Florida.

### Stimuli

Visual and auditory stimuli were created in MATLAB (MathWorks) with the Psychophysics Toolbox (Brainard, 1997; Kleiner et al., 2007) on a PC running Linux Ubuntu 18.04. Visual stimuli were presented on an LCD display (Display ++, Cambridge Research Systems) running at 120 Hz. Visual stimuli were high contrast Gabor patches of moderate spatial frequency (1.18 cycles/°) presented against a black background (0.01 cd/m2). Gabors appeared individually at one of five equally spaced locations. One location was centered at the horizontal meridian on the right side of the screen, and adjacent locations were each separated by an inner angle of 72° (360/5) resulting in an arc length of 8.86 π/5 (~278.3 mm) or 13.23° of visual angle between Gabor centers. The spatial position at which Gabor patches appeared in each trial was quasi-random within each subset of 15 trials to ensure a relatively uniform exposure to the CS+/US contingency over the course of the acquisition trial block. Within every 15-trial subset a Gabor was shown at each of the five spatial positions exactly three times. Each Gabor patch subtended 7.07° of visual angle and had a maximum luminance of 96 cd/m2. The US was a 90 dB sound pressure level (SPL) white noise delivered binaurally through computer speakers located behind participants’ seated location.

### Procedure

Participants were seated in a dimly lit (~60 cd/m2) room with their heads placed on a chin rest to minimize head movements. Viewing distance to the display monitor was set to 120 cm, and distance to the eye-tracker lens was set to 60 cm. Participants were instructed to try to limit blinking to the intertrial interval and to maintain gaze at the central fixation dot throughout testing.

### Data Recording and Analysis

An EyeLink 1000 Plus (SR Research) eye tracker using a 16 mm lens recorded pupil and gaze continuously at 500 Hz. Prior to experimental data collection, the eye tracker was calibrated for each individual. Participants fixated on a nine-point grid, which showed a white circle (5° visual angle) at nine locations against a black background. A second run, following the same 9-point procedure, was then conducted to validate settings obtained in the calibration phase. Pupil diameter was tracked via fitting an ellipse to the thresholded pupil mass, and the illumination level of the infrared signal was set to 100%. Pupil segments outside of a range between 0.01 and 5 mm (~400–16000 in manufacturer arbitrary units) were discarded. Offline, pupil trials were rejected if artifacts such as blinks obscured more than 200 sample points or if they contained any eye movements. Brief (<200 sample points) artifacts were interpolated using piecewise cubic interpolation (see http://dx.doi.org/10.6084/m9.figshare.688002).

#### Pupillometry analysis

The minimum diameter value of the PLR response and the linear slope measuring re-dilation from minimum PLR values to diameter the end of each trial were assessed as dependent variables. Minimum PLR and re-dilation slope were calculated within subjects and for each trial. Slope calculations were computed as the linear slope from the point of minimum pupil diameter when viewing the CS+ stimulus location within a period ranging from 600 ms. to 1200 ms. post-stimulus onset (window determined from inspection of grand-average data, see figure 2) to the pupil diameter at the end of trials prior to US onset (2000 ms. post-stimulus onset). Minimum PLR and slope values were analyzed separately, first with mixed-model Bayesian ANOVAs (Morey & Rouder, 2018) with individuals as random factors, followed by 2 group (CS+ versus CS-) analyses using a Bayesian alternative to traditional *t*-tests (***BEST***: Bayesian Analysis Supersedes the *t*-test; Kruschke, 2013). As measures of pupil activity are known to show substantial inter-individual differences, Cohen’s *d* was calculated as a measure of individual-level effect size for both dependent variables. For these individual effect size measures, standard deviations were obtained from 10,000 iterations of bootstrap resampling of acquisition trials for each participant. Both BEST analyses and computation of individual-level effect sizes were done only for the acquisition trial block, allowing direct comparison to the effect sizes reported in a recent meta-analysis of classical conditioning pupil diameter studies (Finke, Roesmann, Stalder, & Klucken, 2021).

**Figure 1.**
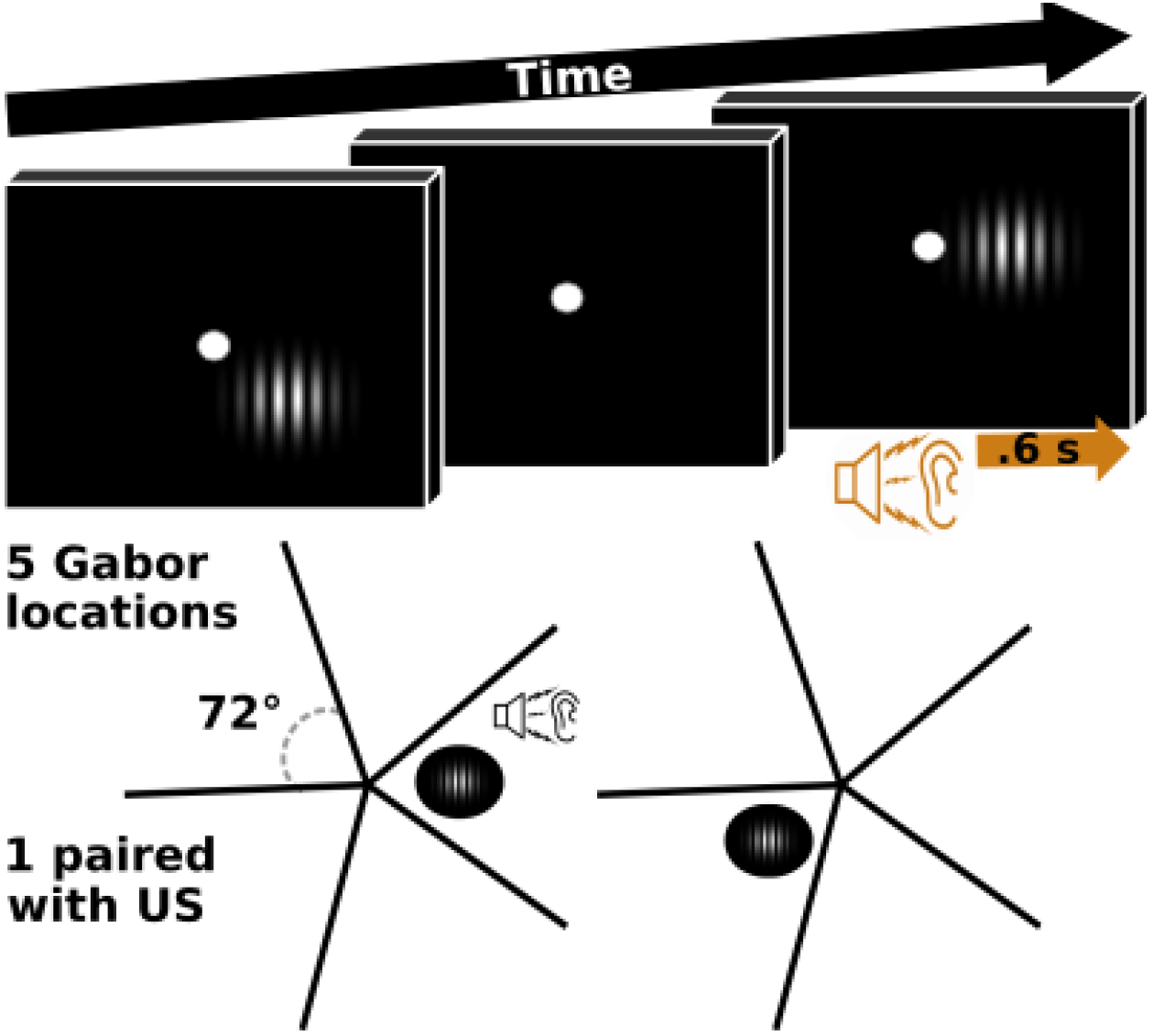
Experimental diagram. An example of 2 acquisition trials, depicting 2 of the possible 5 spatial locations used. Fixation dots are not drawn to scale (enlarged for on-screen/print viewing).

**Figure 2.**
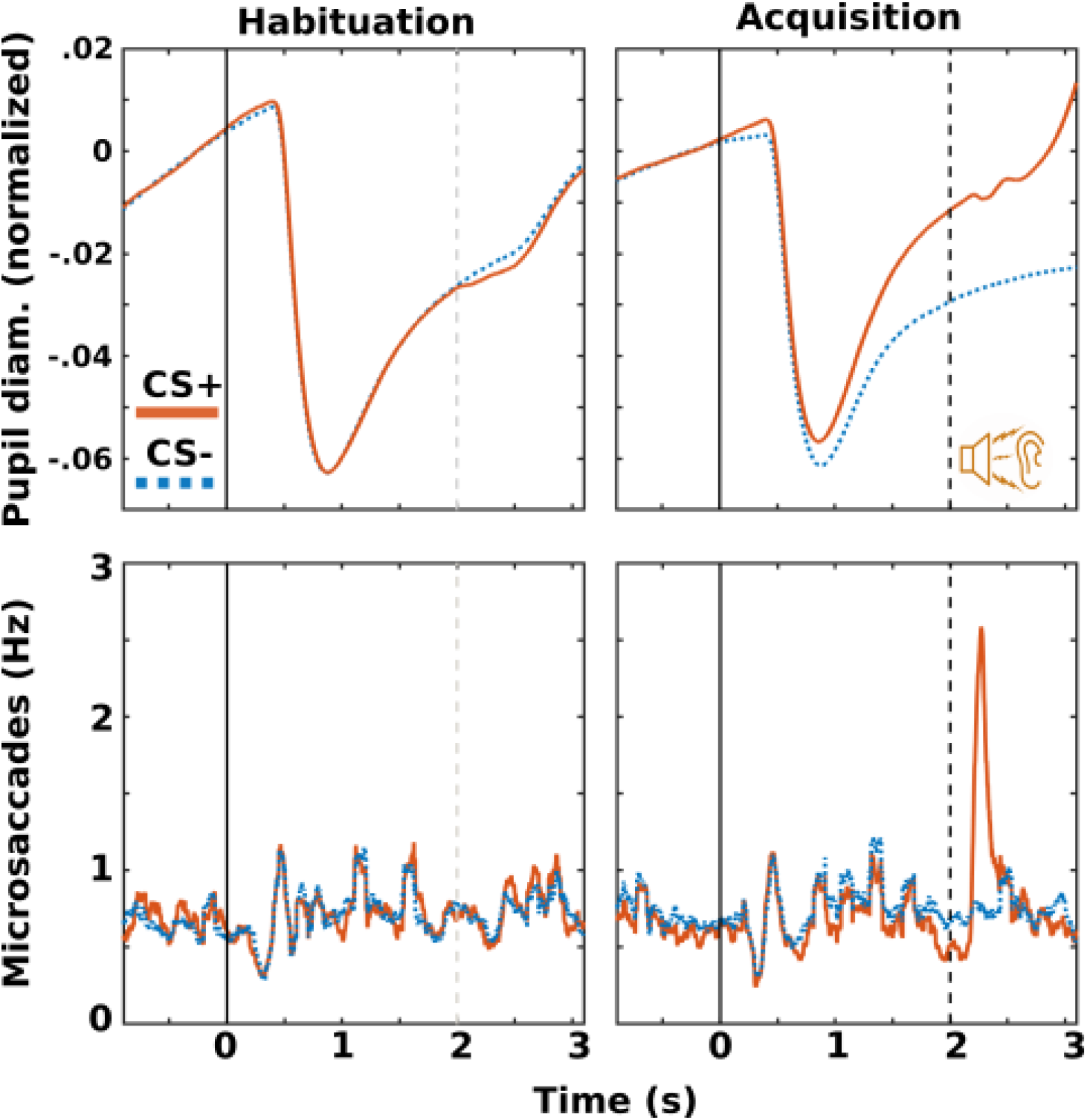
Overall time courses of pupillary responses (top) and microsaccades (bottom) in habituation (left) and acquisition (right) trial blocks. Unconditioned stimulus (US) onset was 2 seconds post-stimulus onset in acquisition trial blocks. Microsaccade time course shown is a 100 ms. moving average.

As previously reported (Friedl & Keil, 2021), pupil diameter for non-aversively associated Gabor patches did not vary as a function of spatial distance from the screen location paired with the US (see supplementary figure 1). Thus, one value for comparison to CS+ responses was obtained by taking the median of the four CS- pupil diameter values within the 600 ms. window defined above. Pupil measures were normalized within individual participants separately for habituation and acquisition trial blocks by dividing each timepoint by the maximum value of the manufacturer (SR research) provided arbitrary units. Baseline pupil diameter was substantially larger in acquisition relative to habituation trial blocks (supplementary figure 2). Due to the typically large end-of-trial differences in pupil diameter between CS- (lower pupil dilation) and CS+ presentations and the quasi-random temporal order (one CS+ trial only rarely followed another CS+ trial) there were clear differences in the acquisition block baseline averages. To reduce CS+/CS- differences due to baseline pupil diameter, the average of baseline pupil diameter in the 400 ms. preceding stimulus onset for each condition was subtracted from each timepoint.

#### Microsaccade analysis

Microsaccades were extracted from artifact-corrected eye position time series data using the algorithm proposed by Engbert and Kliegl (2003; hereafter EK). The EK algorithm was implemented in MATLAB using a modified version of code available from https://github.com/sj971/neurosci-saccade-detection. First, the first derivative was taken over a 6-timepoint sliding window to obtain a velocity time series. Microsaccades were then identified as timepoints in which the Euclidian velocity exceeded 6 standard deviations—a conservative and commonly used threshold (Mihali, van Opheusden, & Ma, 2017)—for at least 12 ms. (Engbert & Kliegl, 2003). The velocity threshold is set on a trial-by-trial basis, ensuring a more conservative threshold on high noise trials. The three user-defined free parameters (size of the sliding window, velocity threshold, and minimum duration exceeding the velocity threshold) were set equal to the values given in Engbert and Kliegl (2003). Data were analyzed using the same software packages described above, with additional time windows of interest detailed in the Results. Unlike pupil diameter, baseline microsaccade rate did not vary meaningfully between CS+ and CS- trials (or between habituation and acquisition blocks) and so no baseline rate adjustment was applied.

#### Mutual information between pupil diameter and microsaccades

Information theory (Shannon & Weaver, 1949) provides the metric of mutual information (MI) to quantify the strength of association between two or more signals. Mutual information measures the reduction in uncertainty in one variable brought about by observing a second variable (Ince, Mazzoni, Petersen, & Panzeri, 2010; Stone, 2018). Mutual information captures any relationship (not only linear relationships) between variables whether those variables are continuous, discrete, or a mixture of the two. In the case of measuring MI between discrete and continuous observations, Ross (2014) provides a method with accompanying MATLAB code that uses a *k*-nearest neighbors estimator, avoiding numerous issues that arise when “binning” continuous data (see Timme & Lapish, 2018). As recommended previously (Kraskov, Stögbauer, & Grassberger, 2004; Ross, 2014), here the user-supplied *k* parameter was set to 3.

## Results

For both minimum PLR value (BF_10_ = 303.34; *F*(1,50) = 9.7) and re-dilation rate (BF_10_ = 1.355 × 10^12^; *F*(1,50) = 19.43), Bayesian ANOVAs including a stimulus type (CS+ versus CS-) by conditioning block (habituation versus acquisition) interaction in addition to main effects and individual participant random factors garnered the strongest support. Results of main interest—the comparison of CS+ and CS- stimuli in the acquisition block—are shown in figure 3. Comparing modulation of the PLR when viewing CS+ versus CS- in the acquisition trial block, 99.8% of the 95% highest density interval (HDI), denoting the most likely values of the posterior distribution given the data and an uninformative prior, lies to the right of zero (BF_10_ 211,338.5; *t*(50) = 5.67). The average effect size (Cohen’s *d*) of PLR modulation at the group level was 0.448 (95% HDI .144 to .746). Re-dilation rate of the pupil was steeper during CS+ trials, with the full HDI of likely difference values falling to the right of zero (BF_10_ = 2.36 × 10^17^; *t*(50) = 9.46). The average effect size of re-dilation rate was 0.826 (95% HDI .367 to 1.32). As shown in figures 4–5, both initial PLR modulation and re-dilation rate showed substantial individual differences in effects. Spearman’s rank order correlation between peak PLR change and re-dilation/unconstriction rate change within participants was –0.106.

**Figure 3.**
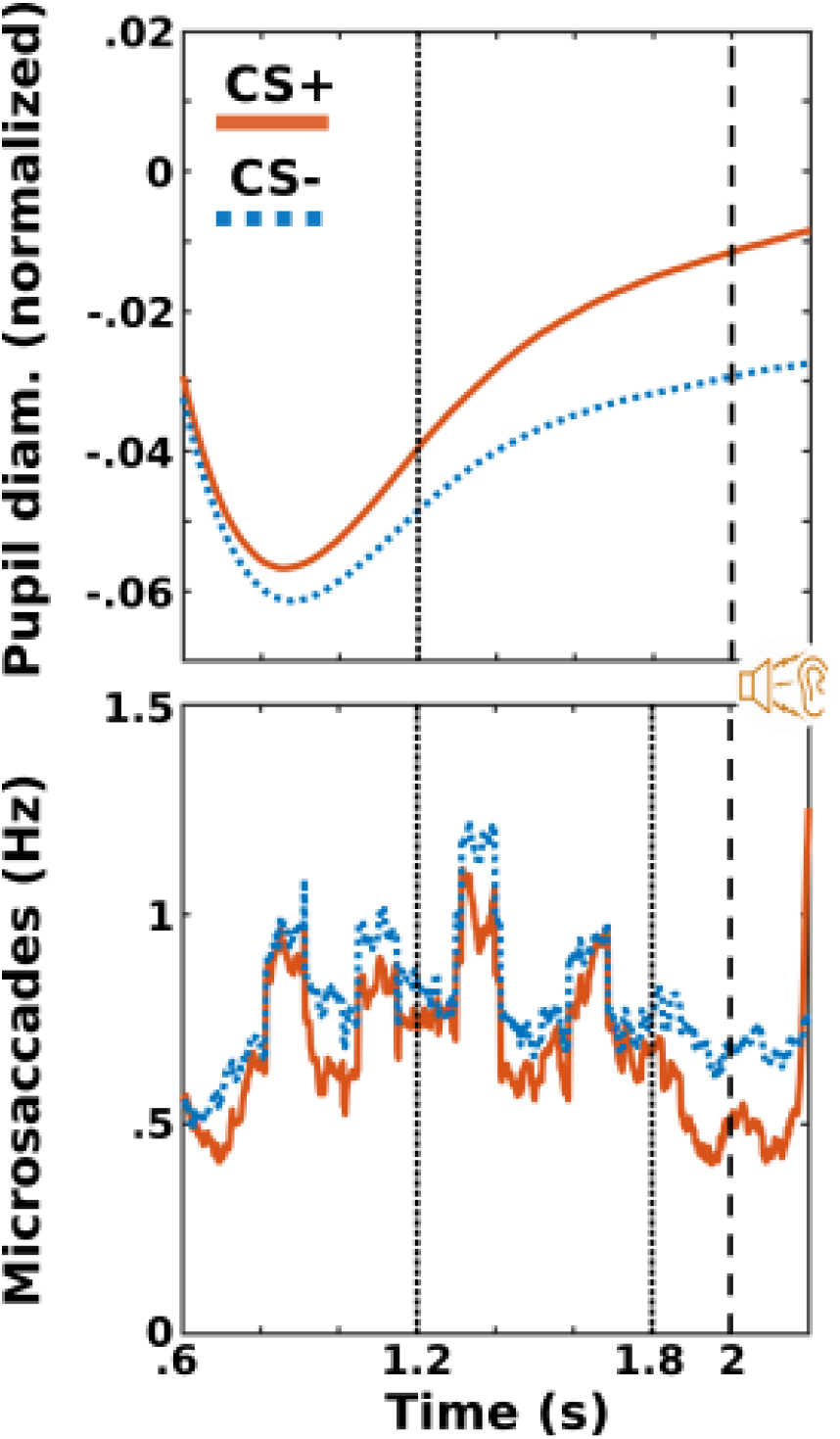
Time windows used in statistical analyses. Pupillary responses (top) and microsaccades (bottom) in acquisition trial blocks. Microsaccade time course shown is a 100 ms. moving average.

**Figure 4.**
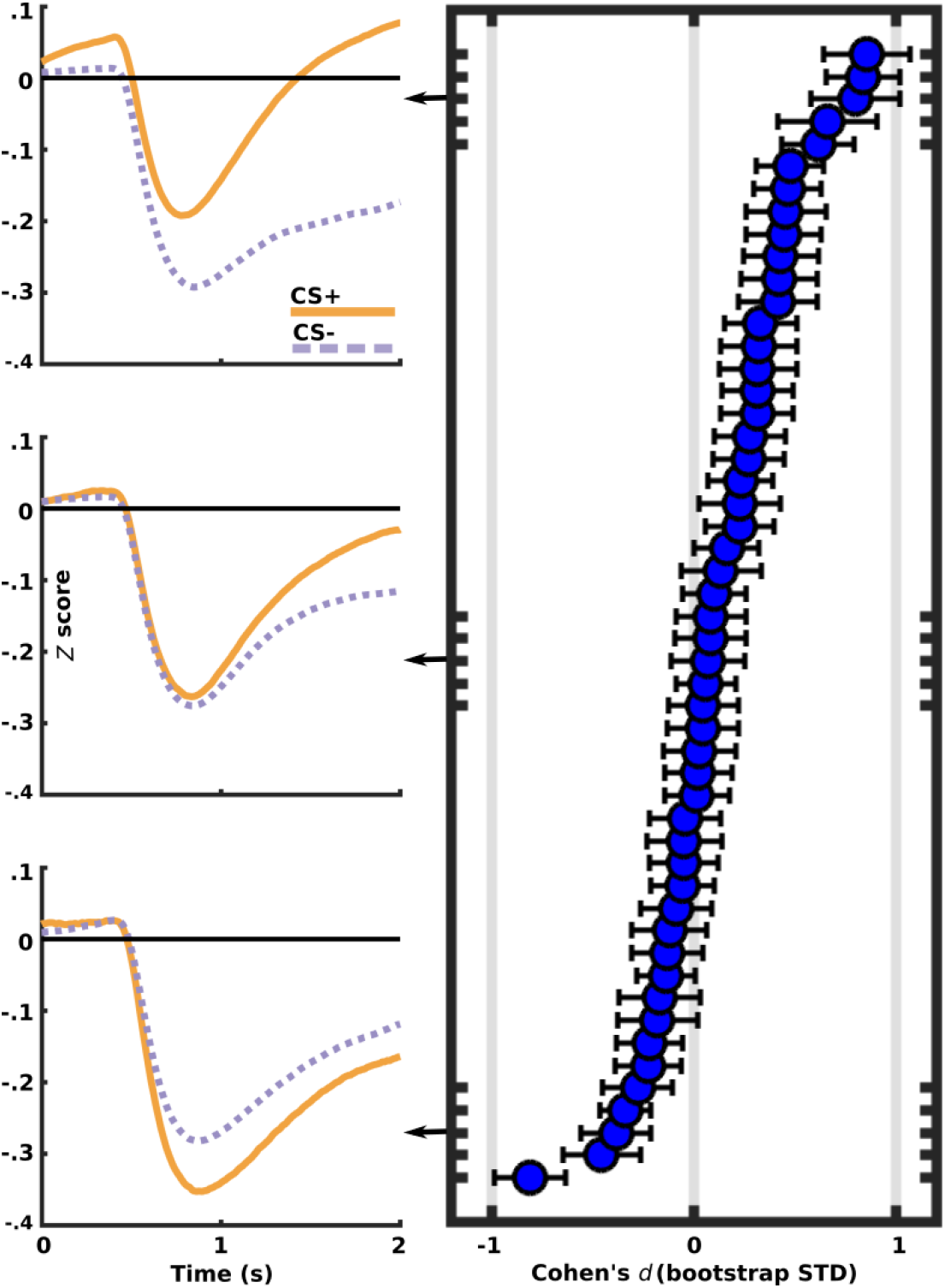
Individual pupillary light response (PLR) effect sizes for CS+ compared to CS- presentations in the acquisition trial block. Positive effect sizes indicate that initial pupil dilation was attenuated for CS+ compared to CS- trials. Figures to the left show time courses for 5-participant sub-sets of the ordered PLR effect.

**Figure 5.**
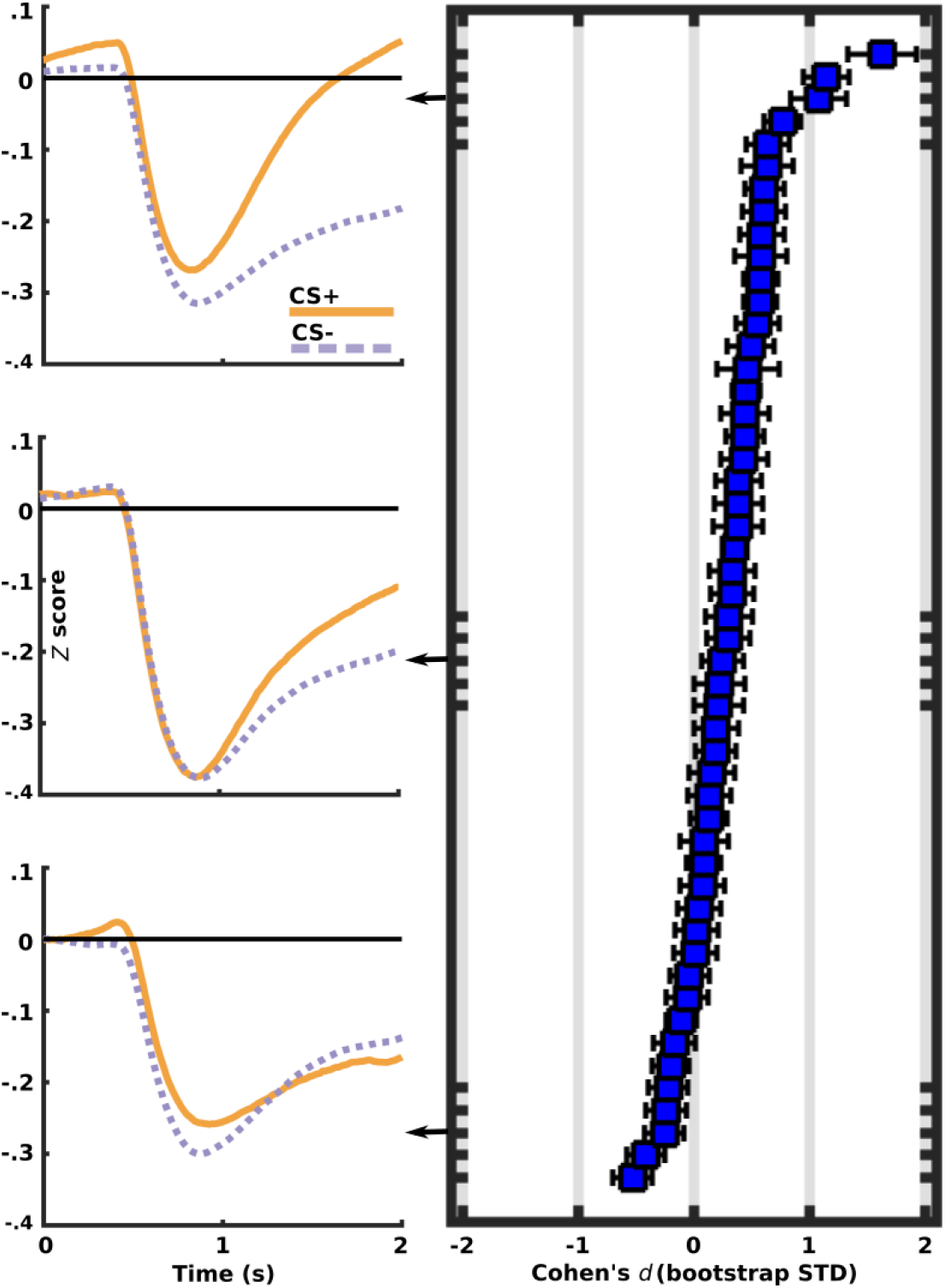
Individual re-dilation/un-constriction rate (slope) effect sizes for CS+ compared to CS- presentations in the acquisition trial block. Positive effect sizes indicate that pupil re-dilation following the PLR was more rapid for CS+ compared to CS- trials. Figures to the left show time courses for 5-participant sub-sets of the ordered re-dilation rate effect.

One time window of microsaccade analysis (600 ms. to 1200 ms. post-stimulus onset) was chosen to overlap with the time window of PLR analysis reported above. Within this time window, group-level differences in the rate of microsaccades were modestly suppressed when viewing a CS+ compared to the (median of the 4 remaining positions) CS- (BF_10_ 11.90; *t*(50) = −3.16). The 95% HDI was entirely below 0 (−.15 to −.03), and the average effect size was −0.434 (95% HDI −.73 to −.13). Two other time windows were examined based upon inspection of the averaged data. Immediately preceding the 600 ms. coinciding with the PLR (i.e. 0 ms. to 600 ms. post stimulus-onset) and the 200 ms. preceding US onset (1,800 ms. to 2,000 ms. post stimulus-onset). From 0-600 ms. group-level differences in the rate of microsaccades were negligible (BF_10_ 0.17; *t*(50) = −0.42). The 95% HDI was nearly centered upon 0, and the average effect size was −0.083 (95% HDI −.385 to .223). In the 200 ms. preceding US onset, microsaccade rate was again lower for the CS+ compared to the CS- (BF_10_ = 825.17; *t*(50) = −4.65). The 95% HDI was entirely below 0 (−.27 to −.11), and the average effect size was −0.71 (95% HDI −1.06 to −.36). The present experimental design did not provide a sufficient number of trials to meaningfully compare microsaccade rates within individuals.

Mutual information between pupil diameter and microsaccade rate over the time windows of 900-1200 ms. and 1,800 ms. to 2,000 post stimulus onset was also assessed. At 900-1200 ms., MI for both the CS+ (BF_10_ = 1.25 × 10^16^) and the CS- (BF_10_ = 1.01 × 10^16^) differed from 0, but with little difference between the signals (*m*_diff_=.0002, BF_10_ =.152). The same pattern of results was observed for the time window immediately preceding US onset: MI was non-zero for both the CS+ (BF_10_ = 1.25 × 10^16^) and CS- (BF_10_ = 1.07 × 10^16^), while the signals did not appreciably differ by conditioning group (*m*_diff_=.0002, BF_10_ =.151).

## Discussion

To elucidate the impacts of learning to associate visual stimuli with unpleasant outcomes at the earliest juncture of perception, we assessed pupillary responses, microsaccade rates, and the mutual information shared by the two measures. Both reflexive dilation of the pupil and the rate of microsaccades following onset of a visual stimulus were found to be modulated by classical conditioning. Shortly following the onset of a conditioned stimulus, illumination-triggered constriction of the pupil was attenuated while the typical suppression of microsaccade rate was more pronounced. Conditioning did not, however, alter the level of statistical association (MI) between the signals. Results suggest that aversive associative learning alters responses in neural regions such as the intermediate layer of the superior colliculus (SCi) common to both pupil dilation and microsaccades.

Pupillometry is becoming an increasingly popular index of differential classical conditioning in human subjects (Finke, Roesmann, Stalder, & Klucken, 2021; Ojala & Bach, 2020; Reinhard & Lachnit, 2002). Despite desirable characteristics such as resistance to habituation (Finke, Roesmann, Stalder, & Klucken, 2021), mixed findings when using pupil diameter as a dependent measure of conditioning have been noted (e.g. Reinhard & Lachnit, 2002; Voigt, 1968). Figures 4–5 (left side), showing subsets of 5 participant averages taken from individuals with low, average, and large effect sizes (within our sample of 51) for conditioned relative to unconditioned stimuli, demonstrate that not only the size, but also the overall direction of effect, could easily differ within smaller samples. Employing an auditory US generally produces smaller changes in pupil diameter compared to using electrical shock (Finke, Roesmann, Stalder, & Klucken, 2021); nevertheless, the average modulation of initial pupillary constriction observed here is in-line with Cohen’s (1992) heuristic for a medium-sized effect (ranging from small to large within individuals). Even slight changes in the relative dilation of the pupil in response to light may confer tremendous perceptual advantages in, for example, times of transitioning light levels such as dawn or dusk (Barlow, 1972, p.3) by altering the trade-off between visual acuity and sensitivity.

Differences in pupil diameter readily distinguish CS+ from CS- stimuli (Ojala & Bach, 2020), but generally prove to be a non-specific indicator of sympathetic system activity or arousal (Bradley, Miccoli, Escrig, & Lang, 2008; Strauch, Greiter, & Huckauf, 2018). Concurrently assessing additional dependent measures such as microsaccads has been proposed as a means of better characterizing changes in the underlying generative process (Strauch, Greiter, & Huckauf, 2018). Here, the rate of change from the peak PLR to US onset was assessed in addition to pupil diameter and microsaccades. The rate of pupil re-dilation/unconstriction following the PLR was much more rapid for CS+ compared to CS- stimulus presentations, and this slope value was only weakly correlated with the degree of PLR modulation within participants. Beyond an early study (Reeves, 1918) reporting that pupil constriction is more rapid than dilation, the rate of pupil change is seldom (if ever) reported. Given the medium-to-large effect sizes observed, the ease of measurement (a simple linear slope was used here), and the near within-subject independence of the measure from pupil diameter, dilation rate could be a promising measure of individual differences in conditioning and other experimental contexts.

Microsaccade rates observed here accord with the average spontaneous rate of 1-2 Hz (Engbert & Kliegl, 2003; Valsecchi & Turatto, 2009) and exhibit the typical post-stimulus biphasic pattern of early inhibition followed by rebound above baseline (Engbert & Kliegl, 2003; Kashihara, Okanoya, & Kawai, 2014; Valsecchi & Turatto, 2009; Yuval-Greenberg, Merriam, & Heeger, 2014). Short latency microsaccade rate suppression is thought to be part of an orienting response (Corneil, & Munoz, 2014; Wang, Blohm, Huang, Boehnke, & Munoz, 2017; Wang, Huang, Brien, & Munoz, 2020), with microsaccades following this period of suppression perhaps serving to eliminate afterimages (Kashihara, 2020). The rebound in microsaccade rate following initial suppression has been found to be impaired when viewing emotional versus neutral natural images (Kashihara, Okanoya, & Kawai, 2014) as well when either viewing (Valsecchi, Betta, & Turatto, 2007) or hearing (Valsecchi & Turatto, 2009) infrequent (oddball) compared to frequent (standard) stimuli. The present investigation found modestly enhanced initial microsaccade rate suppression, but no evidence of an alteration in rebound rate for CS+ compared to CS- visual presentations (figure 3, bottom).

While spontaneous microsaccades prevent visual fading, they are suppressed when performing tasks requiring high visual acuity, such as threading a needle, where a stable retinal image is beneficial (Otero-Millan, Macknik, Serrano-Pedraza, & Martinez-Conde, 2008; Winterson & Collewun, 1976). Initial microsaccade suppression following the appearance of aversively associated CS+ stimuli may therefore be part of an adaptive oculomotor “freezing” response (Badde, Myers, Yuval-Greenberg, & Carrasco, 2020; Rösler & Gamer, 2019) to more accurately process threat-predictive visual cues. On the other hand, although only one visual location in this study predicted an upcoming aversive US, the four remaining screen locations all reliably indicated no upcoming US noise (i.e. they became safety cues). Thus, while only the CS+ location was associated with the US, correctly perceiving *any* screen location reduces the uncertainty about an upcoming US noise equally. The dichotomous discrimination called for here can be likened to a coin flip, where being informed of the outcome of either heads (CS+) or tails (CS-) supplies the same amount of information (1 bit): In either case, uncertainty as to whether or not a US noise is soon to follow is entirely eliminated. From this information-processing perspective, both the CS+ and the CS- presentation locations are relevant to the task of alerting observers to the upcoming state of the world. One possibility is that only the spatial representation of the CS+ location is altered in a neural priority map (Bisley & Mirpour, 2019; Fecteau & Munoz, 2006; Serences & Yantis, 2006) of the visual environment within individuals as a result of the conditioning experience. Such a priority map is thought to be instantiated in SC (Bayguinov, Ghitani, Jackson, & Basso, 2015; Bisley & Mirpour, 2019; White et al., 2017; White, Kan, Levy, Itti, & Munoz, 2017) as well as V1 (Bisley & Mirpour, 2019; Zhang, Zhaoping, Zhou, & Fang, 2012; Zhaoping, 2019). Future work could employ partially occluded stimuli and/or stimuli that are differentially informative of an upcoming US to clarify the extent to which microsaccade rates reflect an attempt to resolve perceptual uncertainty versus altered salience.

Microsaccade rates were also suppressed for the CS+ towards the end of trials, just prior to US onset. A secondary period of suppression has previously been noted in anticipation of upcoming trials (Badde, Myers, Yuval-Greenberg, & Carrasco, 2020; Gao, Yan, & Sun, 2015; Rolfs, Kliegl, & Engbert, 2008). As pupil dilation and saccades are hypothesized to share a common neural origin (Joshi & Gold, 2020; Wang & Munoz, 2015; 2021), late trial microsaccade rate suppression when the pupil is relatively dilated is somewhat surprising. However, sub-threshold stimulation of the SC can cause pupil dilation without triggering eye movements (Joshi, Li, Kalwani, & Gold, 2016; Wang, Boehnke, White, & Munoz, 2012). Further demonstrating that these processes can be dissociated, luminance changes affect pupil diameter but not eye movements (Wang & Munoz, 2021). Different underlying cortical processes could be driving responses in a time-dependent manner within trials, with rapid sub-cortical processing transitioning to higher cognitive processes at later phases (Engbert, 2006). Activity of the locus coeruleus–norepinephrine system (LC-NE), commonly gauged by pupil dilation (Aston-Jones & Cohen, 2005; Joshi, Li, Kalwani, & Gold, 2016; Koss, 1986; Murphy, Robertson, Balsters, & O’connell, 2011), could also contribute to the observed time courses of pupil and microsaccade activity. Norepinephrine can have excitatory or inhibitory effects on target neurons (Aston-Jones & Cohen, 2005; Servan-Schreiber, Printz, & Cohen, 1990) manifesting in cortical layer and dose-dependent fashion throughout the visual system (reviewed in McBurney-Lin, Lu, Zuo, & Yang, 2019). Efforts by participants to maintain central fixation by inhibiting the natural startle reflex to an impending white-noise auditory blast may also have contributed to the depressed pre-US microsaccade rate.

In order for responses to threat-related stimuli learned through classical conditioning to be adaptive in an evolutionary sense, they must help prepare an organism to better deal with its environment (Fanselow & Wassum, 2016; Ramsay, & Woods, 2016). The conjectures ventured above—that microsaccade inhibition could be linked to stabilizing behaviorally-relevant stimuli for better perceptual processing, changes in a neural priority map, and/or to oculomotor system inhibition—are straightforward. On the other hand, postulating potential advantages of a more dilated pupil during CS+ trials in the present experimental context is more challenging. Whether a relatively dilated pupil helps or hinders visual detection or discrimination performance is highly dependent upon experimental conditions (Ajasse, Benosman, & Lorenceau, 2018; Bombeke, Duthoo, Mueller, Hopf, & Boehler, 2016; Mathôt, 2018). In general, a dilated pupil aids detection by increasing photon capture at the retina, while a more constricted pupil enhances the quality of the retinal image, improving discrimination (Binda, Pereverzeva, & Murray, 2013). As there was no task in the employed experimental paradigm, relative pupil dilation when viewing a CS+ may simply reflect increased levels of arousal and/or attention. Consistent with this interpretation, increased phasic NE release has been observed both for target compared to distractor stimuli (Aston-Jones, Rajkowski, & Cohen, 2000; Rajkowski, Majczynski, Clayton, & Aston-Jones, 2004) as well as to conditioned stimuli (Bouret & Richmond, 2009). Correspondingly, there is evidence that arousal biases processing resources towards task-relevant relative to non-relevant stimuli (Mather & Sutherland, 2011), and that such bias increases pupil diameter (Mather, Clewett, Sakaki, & Harley, 2016).

The processes of arousal and attention are sometimes held to be synonymous (e.g. Dayan, Kakade, & Montague, 2000) and sometimes to be distinct but overlapping phenomena (e.g. Coull, 1998; Sadaghiani & Kleinschmidt, 2016). The present work focuses on arousal, given the linkage of this term to studies of pupillary dilation and LC-NE system function. However, as no efforts were made to disentangle these closely related concepts, no strong statement favoring either attention or arousal-based theories can be made. There is considerable overlap between “attention networks” and the brain networks that drive both eye movements (Büchel et al., 1998; Corbetta et al., 1998; Parr & Friston, 2017; 2019) and pupil dilation (Alnæs et al., 2014). As mentioned in the introduction, both microsaccades and pupil diameter changes have also been shown to track shifts in covert attention. Future work would be needed to resolve this arousal/attention definitional ambiguity. Pupil frequency tagging (PFT), described by Naber, Alvarez, and Nakayama (2013), is one method of isolating the impact of spatial attention on pupillary responses that could prove useful in such an endeavor.

Compared with brain regions constituting the traditional “fear network”, the effects of classical conditioning on sensory processing areas remain poorly understood. This work adds to a body of evidence suggesting early sensory systems are important targets of neural adaptation triggered by experienced aversive associations: In addition to visual (e.g. Friedl & Keil, 2020; 2021; McTeague, Gruss, & Keil, 2015; Santos-Mayo, de Echegaray, & Moratti, 2022), auditory (e.g. Letzkus et al. 2011; Quirk, Armony, & LeDoux, 1997; Wood, Angeloni, Oxman, Clopath, & Geffen, 2022), and olfactory (You, Novak, Clancy, & Li, 2022) cortices, subcortical networks that drive eye movements and/or pupillary responses are implicated here. Another pressing gap in the current classical conditioning literature base concerns individual-level effects of experimental manipulations (Beckers, Krypotos, Boddez, Effting, & Kindt, 2013; Lonsdorf & Merz, 2017). Ample evidence for the existence and direction of group-level conditioning effects on pupillary responses exist: Conditioned stimuli prompt a relatively dilated/unconstricted pupil. Individual heterogeneity in initial PLR modulation and rate of pupil re-dilation, as demonstrated here, highlight these easily assessable measures as promising indicators of individual-level differences in autonomic reactivity. As suggested previously (Strauch, Greiter, & Huckauf, 2018), microsaccade rates in the present experimental context generally support inferences derived from pupillary measures while showing smaller effect sizes. Microsaccade rates thus appear to be more suitable for group-level analyses, and less promising as a means of probing individual differences in associative learning.

## Supplementary Information

Overall cortical arousal levels tend to increase during acquisition trials compared to habitation trials (Lissek et al., 2008). Reflecting this, baseline pupil diameter was substantially greater in the acquisition trial block (supplementary figure 2, top inset). Baseline pupil diameter has been linked to tonic activity of NE (Aston-Jones & Cohen, 2005) as well as information processing (Poock, 1973; Zénon, 2019). Entropy quantifies the average amount of Shannon information in a distribution (Stone, 2018, p.13), and is applied here to the normalized pupil diameter values in the pre-stimulus period at varying points of the experiment to gauge changes in information processing. Entropy of the distribution of baseline pupil diameter values fluctuates throughout the experimental session such that periods of a more dilated/less constricted pupil is associated with states of lower entropy/uncertainty.

---

**Supplemental Figure 1.**
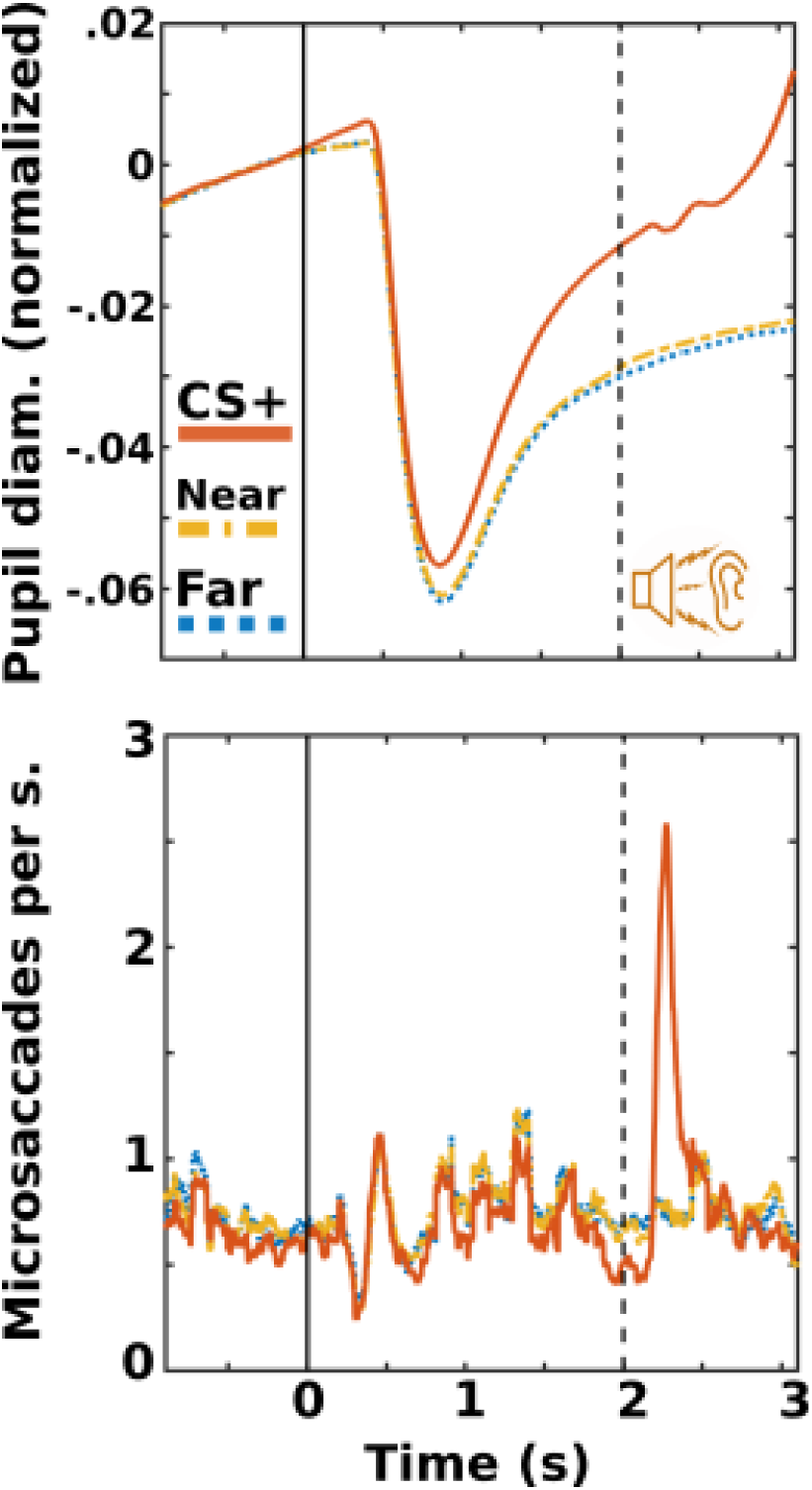
Overall average timecourse for pupillary responses (top) and microsaccade rate (bottom) as a function of distance from the CS+ screen position in the acquisition trial block. As both measures showed little differentiation between the near and far CS- locations, all main figures and comparison statistics combined near and far responses into one aggregate CS- value (see Method section for details). Microsaccade time course shown is a 100 ms. moving average.

**Supplemental Figure 2.**
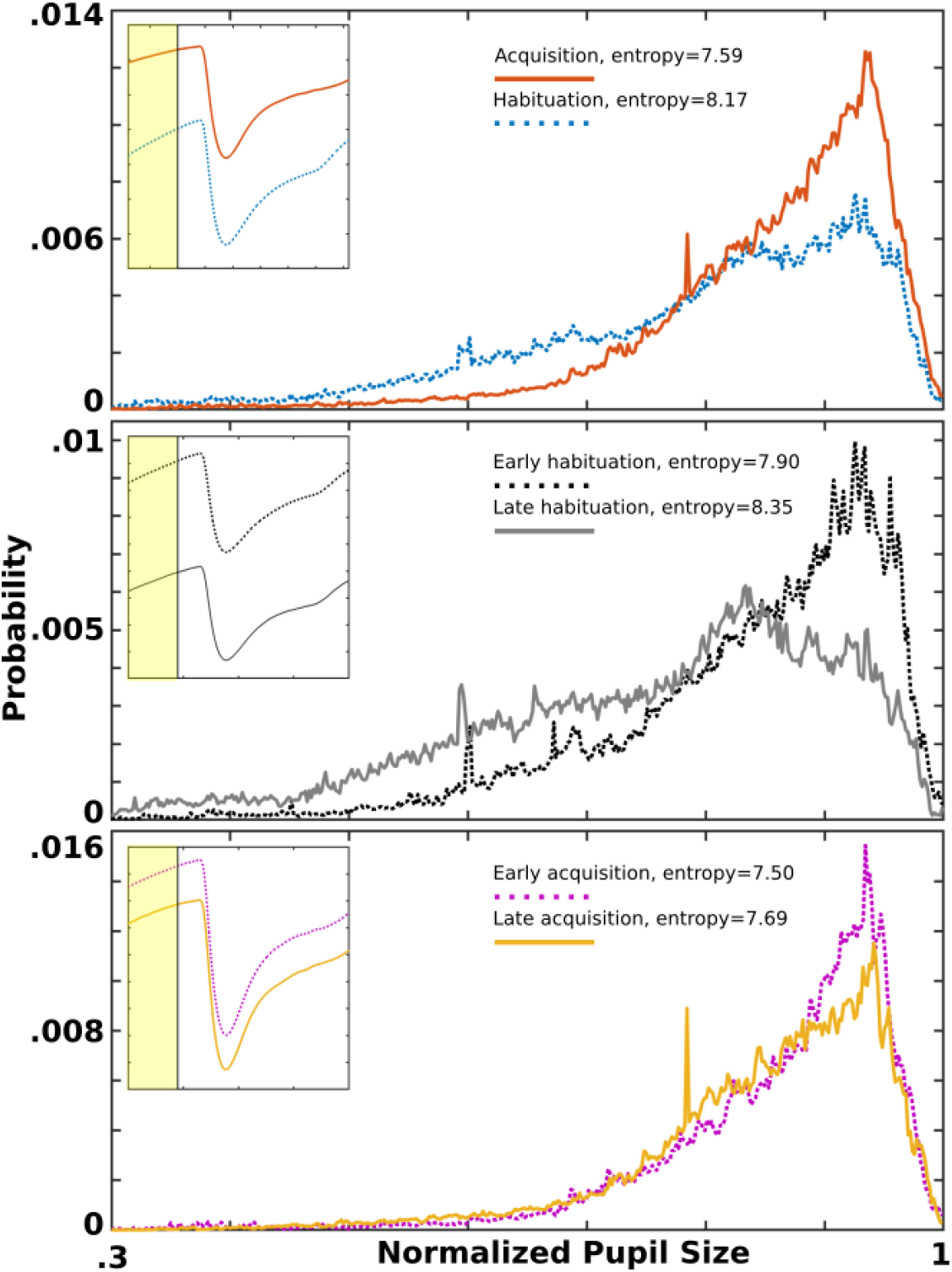
Entropy in overall probability distributions of pupil diameter values in pre-stimulus baseline period (yellow shaded area of insets). Acquisition and habituation trial blocks (top). First half (trials 1-75) and second half (trials 76-150) of habituation (middle). First half (trials 151-250) and second half (trials 251-350) of acquisition (bottom).

## Notes

https://github.com/sj971/neurosci-saccade-detection

http://dx.doi.org/10.6084/m9.figshare.688002

### Competing Interest Statement

The authors have declared no competing interest.

## References

Ajasse, S., Benosman, R. B., & Lorenceau, J. (2018). Effects of pupillary responses to luminance and attention on visual spatial discrimination. Journal of vision, 18(11), 6–6.

Akdoğan, B., Balci, F., & van Rijn, H. (2016). Temporal expectation indexed by pupillary response. Timing & Time Perception, 4(4), 354–370.

Alnæs, D., Sneve, M. H., Espeseth, T., Endestad, T., van de Pavert, S. H. P., & Laeng, B. (2014). Pupil size signals mental effort deployed during multiple object tracking and predicts brain activity in the dorsal attention network and the locus coeruleus. Journal of vision, 14(4), 1–1.

Aston-Jones, G., & Cohen, J. D. (2005). An integrative theory of locus coeruleus-norepinephrine function: adaptive gain and optimal performance. Annu. Rev. Neurosci., 28, 403–450.

Aston-Jones, G., Rajkowski, J., & Cohen, J. (2000). Locus coeruleus and regulation of behavioral flexibility and attention. Progress in brain research, 126, 165–182.

Badde, S., Myers, C. F., Yuval-Greenberg, S., & Carrasco, M. (2020). Oculomotor freezing reflects tactile temporal expectation and aids tactile perception. Nature communications, 11(1), 1–9.

Barlow, H. B. (1972). Dark and light adaptation: psychophysics. In Visual psychophysics (pp. 1–28). Springer, Berlin, Heidelberg.

Bayguinov, P. O., Ghitani, N., Jackson, M. B., & Basso, M. A. (2015). A hard-wired priority map in the superior colliculus shaped by asymmetric inhibitory circuitry. Journal of neurophysiology, 114(1), 662–676.

Beckers, T., Krypotos, A. M., Boddez, Y., Effting, M., & Kindt, M. (2013). What’s wrong with fear conditioning?. Biological psychology, 92(1), 90–96.

Benedetto, A., & Binda, P. (2016). Dissociable saccadic suppression of pupillary and perceptual responses to light. Journal of neurophysiology, 115(3), 1243–1251.

Betta, E., Galfano, G., & Turatto, M. (2007). Microsaccadic response during inhibition of return in a target–target paradigm. Vision Research, 47(3), 428–436.

Binda, P., Pereverzeva, M., & Murray, S. O. (2013). Attention to bright surfaces enhances the pupillary light reflex. Journal of Neuroscience, 33(5), 2199–2204.

Binda, P., Pereverzeva, M., & Murray, S. O. (2013). Pupil constrictions to photographs of the sun. Journal of Vision, 13(6), 8–8.

Bisley, J. W., & Mirpour, K. (2019). The neural instantiation of a priority map. Current opinion in psychology, 29, 108–112.

Blanchard, D. C., & Blanchard, R. J. (1972). Innate and conditioned reactions to threat in rats with amygdaloid lesions. Journal of comparative and physiological psychology, 81(2), 281.

Bombeke, K., Duthoo, W., Mueller, S. C., Hopf, J. M., & Boehler, C. N. (2016). Pupil size directly modulates the feedforward response in human primary visual cortex independently of attention. NeuroImage, 127, 67–73.

Bouret, S., & Richmond, B. J. (2009). Relation of locus coeruleus neurons in monkeys to Pavlovian and operant behaviors. Journal of neurophysiology, 101(2), 898–911.

Bradley, M. M. (2009). Natural selective attention: Orienting and emotion. Psychophysiology, 46(1), 1–11.

Bradley, M. M., Miccoli, L., Escrig, M. A., & Lang, P. J. (2008). The pupil as a measure of emotional arousal and autonomic activation. Psychophysiology, 45(4), 602–607.

Bradley, M. M., Sapigao, R. G., & Lang, P. J. (2017). Sympathetic ANS modulation of pupil diameter in emotional scene perception: Effects of hedonic content, brightness, and contrast. Psychophysiology, 54(10), 1419–1435.

Büchel, C., Josephs, O., Rees, G., Turner, R., Frith, C. D., & Friston, K. J. (1998). The functional anatomy of attention to visual motion. A functional MRI study. Brain: a journal of neurology, 121(7), 1281–1294.

Campbell, F. W., & Gregory, A. H. (1960). Effect of size of pupil on visual acuity. Nature, 187(4743), 1121–1123.

Carrasco, M. (2018). How visual spatial attention alters perception. Cognitive processing, 19(1), 77–88.

Chen, C. Y., Ignashchenkova, A., Thier, P., & Hafed, Z. M. (2015). Neuronal response gain enhancement prior to microsaccades. Current Biology, 25(16), 2065–2074.

Cohen, J. (1992). A power primer. Psychological bulletin, 112(1), 155.

Corbetta, M., Akbudak, E., Conturo, T. E., Snyder, A. Z., Ollinger, J. M., Drury, H. A., … & Shulman, G. L. (1998). A common network of functional areas for attention and eye movements. Neuron, 21(4), 761–773.

Corneil, B. D., & Munoz, D. P. (2014). Overt responses during covert orienting. Neuron, 82(6), 1230–1243.

Coull, J. T. (1998). Neural correlates of attention and arousal: insights from electrophysiology, functional neuroimaging and psychopharmacology. Progress in neurobiology, 55(4), 343–361.

Domjan, M. (2018). The essentials of conditioning and learning. American Psychological Association.

Dayan, P., Kakade, S., & Montague, P. R. (2000). Learning and selective attention. Nature neuroscience, 3(11), 1218–1223.

Engbert, R. (2006). Microsaccades: A microcosm for research on oculomotor control, attention, and visual perception. Progress in brain research, 154, 177–192.

Engbert, R., & Kliegl, R. (2003). Microsaccades uncover the orientation of covert attention. Vision research, 43(9), 1035–1045.

Fanselow, M. S., & Wassum, K. M. (2016). The origins and organization of vertebrate Pavlovian conditioning. Cold Spring Harbor perspectives in biology, 8(1), a021717.

Fecteau, J. H., & Munoz, D. P. (2006). Salience, relevance, and firing: a priority map for target selection. Trends in cognitive sciences, 10(8), 382–390.

Finke, J. B., Roesmann, K., Stalder, T., & Klucken, T. (2021). Pupil dilation as an index of Pavlovian conditioning. A systematic review and meta-analysis. Neuroscience & Biobehavioral Reviews, 130, 351–368.

Friedl, W. M., & Keil, A. (2020). Effects of experience on spatial frequency tuning in the visual system: behavioral, visuocortical, and alpha-band responses. Journal of Cognitive Neuroscience, 32(6), 1153–1169.

Friedl, W. M., & Keil, A. (2021). Aversive Conditioning of Spatial Position Sharpens Neural Population-Level Tuning in Visual Cortex and Selectively Alters Alpha-Band Activity. Journal of Neuroscience, 41(26), 5723–5733.

Gao, X., Yan, H., & Sun, H. J. (2015). Modulation of microsaccade rate by task difficulty revealed through between-and within-trial comparisons. Journal of vision, 15(3), 3–3.

Hafed, Z. M., & Clark, J. J. (2002). Microsaccades as an overt measure of covert attention shifts. Vision research, 42(22), 2533–2545.

Hafed, Z. M., & Krauzlis, R. J. (2012). Similarity of superior colliculus involvement in microsaccade and saccade generation. Journal of neurophysiology, 107(7), 1904–1916.

Horowitz, T. S., Fine, E. M., Fencsik, D. E., Yurgenson, S., & Wolfe, J. M. (2007). Fixational eye movements are not an index of covert attention. Psychological science, 18(4), 356–363.

Hudspeth, A. J., Jessell, T. M., Kandel, E. R., Schwartz, J. H., & Siegelbaum, S. A. (Eds.). (2013). Principles of neural science. McGraw-Hill, Health Professions Division.

Galfano, G., Betta, E., & Turatto, M. (2004). Inhibition of return in microsaccades. Experimental Brain Research, 159(3), 400–404.

Ibbotson, M., & Krekelberg, B. (2011). Visual perception and saccadic eye movements. Current opinion in neurobiology, 21(4), 553–558.

Ince, R. A., Mazzoni, A., Petersen, R. S., & Panzeri, S. (2010). Open source tools for the information theoretic analysis of neural data. Frontiers in neuroscience, 3, 11.

Joshi, S., & Gold, J. I. (2020). Pupil size as a window on neural substrates of cognition. Trends in cognitive sciences, 24(6), 466–480.

Joshi, S., Li, Y., Kalwani, R. M., & Gold, J. I. (2016). Relationships between pupil diameter and neuronal activity in the locus coeruleus, colliculi, and cingulate cortex. Neuron, 89(1), 221–234.

Kashihara, K. (2020). Microsaccadic modulation evoked by emotional events. Journal of Physiological Anthropology, 39(1), 1–11.

Kashihara, K., Okanoya, K., & Kawai, N. (2014). Emotional attention modulates microsaccadic rate and direction. Psychological research, 78(2), 166–179.

Koss, M. C. (1986). Pupillary dilation as an index of central nervous system α2-adrenoceptor activation. Journal of pharmacological methods, 15(1), 1–19.

Kraskov, A., Stögbauer, H., & Grassberger, P. (2004). Estimating mutual information. Physical review E, 69(6), 066138.

Kruschke, J. K. (2013). Bayesian estimation supersedes the t test. Journal of Experimental Psychology: General, 142(2), 573.

Lang, P. J., Bradley, M. M., & Cuthbert, B. N. (1997). Motivated attention: Affect, activation, and action. Attention and orienting: Sensory and motivational processes, 97, 135.

Laubrock, J., Engbert, R., Rolfs, M., & Kliegl, R. (2007). Microsaccades are an index of covert attention: commentary on Horowitz, Fine, Fencsik, Yurgenson, and Wolfe (2007). Psychological Science, 18(4), 364–366.

Laughlin, S. B. (1992). Retinal information capacity and the function of the pupil. Ophthalmic and Physiological Optics, 12(2), 161–164.

Letzkus, J. J., Wolff, S. B., Meyer, E. M., Tovote, P., Courtin, J., Herry, C., & Lüthi, A. (2011). A disinhibitory microcircuit for associative fear learning in the auditory cortex. Nature, 480(7377), 331–335.

LeDoux, J. E. (2000). Emotion circuits in the brain. Annual review of neuroscience, 23(1), 155–184.

LeDoux, J. E., Cicchetti, P., Xagoraris, A., & Romanski, L. M. (1990). The lateral amygdaloid nucleus: sensory interface of the amygdala in fear conditioning. Journal of neuroscience, 10(4), 1062–1069.

Lissek, S., Biggs, A. L., Rabin, S. J., Cornwell, B. R., Alvarez, R. P., Pine, D. S., & Grillon, C. (2008). Generalization of conditioned fear-potentiated startle in humans: experimental validation and clinical relevance. Behaviour research and therapy, 46(5), 678–687.

Liversedge, S. P., & Findlay, J. M. (2000). Saccadic eye movements and cognition. Trends in cognitive sciences, 4(1), 6–14.

Lonsdorf, T. B., & Merz, C. J. (2017). More than just noise: Inter-individual differences in fear acquisition, extinction and return of fear in humans-Biological, experiential, temperamental factors, and methodological pitfalls. Neuroscience & Biobehavioral Reviews, 80, 703–728.

Lorber, M., Zuber, B. L., & Stark, L. (1965). Suppression of the pupillary light reflex in binocular rivalry and saccadic suppression. Nature, 208(5010), 558–560.

Mackintosh, N. J. (1975). A theory of attention: Variations in the associability of stimuli with reinforcement. Psychological review, 82(4), 276.

Martinez-Conde, S., Macknik, S. L., & Hubel, D. H. (2004). The role of fixational eye movements in visual perception. Nature reviews neuroscience, 5(3), 229–240.

Mather, M., Clewett, D., Sakaki, M., & Harley, C. W. (2016). Norepinephrine ignites local hotspots of neuronal excitation: How arousal amplifies selectivity in perception and memory. Behavioral and Brain Sciences, 39.

Mather, M., & Sutherland, M. R. (2011). Arousal-biased competition in perception and memory. Perspectives on psychological science, 6(2), 114–133.

Mathôt, S. (2018). Pupillometry: psychology, physiology, and function. Journal of Cognition, 1(1).

Mathôt, S., & Ivanov, Y. (2019). The effect of pupil size and peripheral brightness on detection and discrimination performance. PeerJ, 7, e8220.

Mathôt, S., Van der Linden, L., Grainger, J., & Vitu, F. (2013). The pupillary light response reveals the focus of covert visual attention. PloS one, 8(10), e78168.

Martinez-Conde, S., Macknik, S. L., Troncoso, X. G., & Dyar, T. A. (2006). Microsaccades counteract visual fading during fixation. Neuron, 49(2), 297–305.

McBurney-Lin, J., Lu, J., Zuo, Y., & Yang, H. (2019). Locus coeruleus-norepinephrine modulation of sensory processing and perception: A focused review. Neuroscience & Biobehavioral Reviews, 105, 190–199.

McTeague, L. M., Gruss, L., & Keil, A. (2015). Aversive learning shapes neuronal orientation tuning in human visual cortex. Nature communications, 6(1), 1–8.

Mihali, A., van Opheusden, B., & Ma, W. J. (2017). Bayesian microsaccade detection. Journal of vision, 17(1), 13–13.

Morey, R.D., & Rouder, J.N. (2018). BayesFactor: Computation of Bayes Factors for Common Designs. R package version 0.9.12-4.2. https://CRAN.R-project.org/package=BayesFactor

Murphy, P. R., Robertson, I. H., Balsters, J. H., & O’connell, R. G. (2011). Pupillometry and P3 index the locus coeruleus–noradrenergic arousal function in humans. Psychophysiology, 48(11), 1532–1543.

Naber, M., Alvarez, G. A., & Nakayama, K. (2013). Tracking the allocation of attention using human pupillary oscillations. Frontiers in psychology, 4, 919.

Najemnik, J., & Geisler, W. S. (2005). Optimal eye movement strategies in visual search. Nature, 434(7031), 387–391.

Ojala, K. E., & Bach, D. R. (2020). Measuring learning in human classical threat conditioning: translational, cognitive and methodological considerations. Neuroscience & Biobehavioral Reviews, 114, 96–112.

Otero-Millan, J., Troncoso, X. G., Macknik, S. L., Serrano-Pedraza, I., & Martinez-Conde, S. (2008). Saccades and microsaccades during visual fixation, exploration, and search: foundations for a common saccadic generator. Journal of vision, 8(14), 21–21.

Pandey, P., & Ray, S. (2021). Pupil dynamics: A potential proxy of neural preparation for goal-directed eye-movement. European Journal of Neuroscience.

Panitz, C., Keil, A., & Mueller, E. M. (2019). Extinction-resistant attention to long-term conditioned threat is indexed by selective visuocortical alpha suppression in humans. Scientific reports, 9(1), 1–9.

Parr, T., & Friston, K. J. (2017). The active construction of the visual world. Neuropsychologia, 104, 92–101.

Parr, T., & Friston, K. J. (2019). Attention or salience?. Current opinion in psychology, 29, 1–5.

Peinkhofer, C., Knudsen, G. M., Moretti, R., & Kondziella, D. (2019). Cortical modulation of pupillary function: systematic review. PeerJ, 7, e6882.

Poock, G. K. (1973). Information processing vs pupil diameter. Perceptual and motor skills, 37(3), 1000–1002.

Privitera, C. M., Carney, T., Klein, S., & Aguilar, M. (2014). Analysis of microsaccades and pupil dilation reveals a common decisional origin during visual search. Vision Research, 95, 43–50.

Quirk, G. J., Armony, J. L., & LeDoux, J. E. (1997). Fear conditioning enhances different temporal components of tone-evoked spike trains in auditory cortex and lateral amygdala. Neuron, 19(3), 613–624.

Rajkowski, J., Majczynski, H., Clayton, E., & Aston-Jones, G. (2004). Activation of monkey locus coeruleus neurons varies with difficulty and performance in a target detection task. Journal of neurophysiology, 92(1), 361–371.

Ramsay, D. S., & Woods, S. C. (2016). Physiological regulation: how it really works. Cell metabolism, 24(3), 361–364.

Reeves, P. (1918). Rate of pupillary dilation and contraction. Psychological Review, 25(4), 330.

Reinhard, G., & Lachnit, H. (2002). Differential conditioning of anticipatory pupillary dilation responses in humans. Biological psychology, 60(1), 51–68.

Rolfs, M., Kliegl, R., & Engbert, R. (2008). Toward a model of microsaccade generation: The case of microsaccadic inhibition. Journal of vision, 8(11), 5–5.

Rösler, L., & Gamer, M. (2019). Freezing of gaze during action preparation under threat imminence. Scientific reports, 9(1), 1–9.

Ross, B. C. (2014). Mutual information between discrete and continuous data sets. PloS one, 9(2), e87357.

Rucci, M., Iovin, R., Poletti, M., & Santini, F. (2007). Miniature eye movements enhance fine spatial detail. Nature, 447(7146), 852–855.

Sadaghiani, S., & Kleinschmidt, A. (2016). Brain networks and α-oscillations: structural and functional foundations of cognitive control. Trends in cognitive sciences, 20(11), 805–817.

Santos-Mayo, A., de Echegaray, J., & Moratti, S. (2022). Conditioned up and down modulations of short latency gamma band oscillations in visual cortex during fear learning in humans. Scientific reports, 12(1), 1–10.

Serences, J. T., & Yantis, S. (2006). Selective visual attention and perceptual coherence. Trends in cognitive sciences, 10(1), 38–45.

Servan-Schreiber, D., Printz, H., & Cohen, J. D. (1990). A network model of catecholamine effects: gain, signal-to-noise ratio, and behavior. Science, 249(4971), 892–895.

Shannon, C. E., & Weaver, W. (1949). The mathematical theory of information. Urbana: University of Illinois Press, 97.

Sirois, S., & Brisson, J. (2014). Pupillometry. Wiley Interdisciplinary Reviews: Cognitive Science, 5(6), 679–692.

Strauch, C., Greiter, L., & Huckauf, A. (2018). Pupil dilation but not microsaccade rate robustly reveals decision formation. Scientific reports, 8(1), 1–9.

Stone, J. V. (2018). Principles of neural information theory: computational neuroscience and metabolic efficiency (tutorial introductions). Tutorial Introductions.

Thompson, R. F. (1983). Neuronal substrates of simple associative learning: Classical conditioning. Trends in Neurosciences, 6, 270–275.

Timme, N. M., & Lapish, C. (2018). A tutorial for information theory in neuroscience. eneuro, 5(3).

Valsecchi, M., Betta, E., & Turatto, M. (2007). Visual oddballs induce prolonged microsaccadic inhibition. Experimental Brain Research, 177(2), 196–208.

Valsecchi, M., & Turatto, M. (2009). Microsaccadic responses in a bimodal oddball task. Psychological research, 73(1), 23–33.

Voigt, W. H. (1968). Conditioning the human pupillary response. Perceptual and motor skills, 26(3), 975–982.

You, Y., Novak, L. R., Clancy, K. J., & Li, W. (2022). Pattern differentiation and tuning shift in human sensory cortex underlie long-term threat memory. Current Biology.

Yuval-Greenberg, S., Merriam, E. P., & Heeger, D. J. (2014). Spontaneous microsaccades reflect shifts in covert attention. Journal of Neuroscience, 34(41), 13693–13700.

Wang, C. A., Boehnke, S. E., White, B. J., & Munoz, D. P. (2012). Microstimulation of the monkey superior colliculus induces pupil dilation without evoking saccades. Journal of Neuroscience, 32(11), 3629–3636.

Wang, C. A., Huang, J., Brien, D. C., & Munoz, D. P. (2020). Saliency and priority modulation in a pop-out paradigm: pupil size and microsaccades. Biological Psychology, 153, 107901.

Wang, C. A., Blohm, G., Huang, J., Boehnke, S. E., & Munoz, D. P. (2017). Multisensory integration in orienting behavior: Pupil size, microsaccades, and saccades. Biological psychology, 129, 36–44.

Wang, C. A., & Munoz, D. P. (2015). A circuit for pupil orienting responses: implications for cognitive modulation of pupil size. Current opinion in neurobiology, 33, 134–140.

Wang, C. A., & Munoz, D. P. (2021). Coordination of pupil and saccade responses by the superior colliculus. Journal of Cognitive Neuroscience, 33(5), 919–932.

White, B. J., Berg, D. J., Kan, J. Y., Marino, R. A., Itti, L., & Munoz, D. P. (2017). Superior colliculus neurons encode a visual saliency map during free viewing of natural dynamic video. Nature communications, 8(1), 1–9.

White, B. J., Kan, J. Y., Levy, R., Itti, L., & Munoz, D. P. (2017). Superior colliculus encodes visual saliency before the primary visual cortex. Proceedings of the National Academy of Sciences, 114(35), 9451–9456.

Whittaker, S. G., & Cummings, R. W. (1990). Foveating saccades. Vision Research, 30(9), 1363–1366.

Winterson, B. J., & Collewun, H. (1976). Microsaccades during finely guided visuomotor tasks. Vision research, 16(12), 1387–1390.

Wood, K. C., Angeloni, C. F., Oxman, K., Clopath, C., & Geffen, M. N. (2022). Neuronal activity in sensory cortex predicts the specificity of learning in mice. Nature communications, 13(1), 1–15.

Zénon, A. (2019). Eye pupil signals information gain. Proceedings of the Royal Society B, 286(1911), 20191593.

Zhang, X., Zhaoping, L., Zhou, T., & Fang, F. (2012). Neural activities in V1 create a bottom-up saliency map. Neuron, 73(1), 183–192.

Zhaoping, L. (2019). A new framework for understanding vision from the perspective of the primary visual cortex. Current Opinion in Neurobiology, 58, 1–10.

